# A pair of non-competing neutralizing human monoclonal antibodies protecting from disease in a SARS-CoV-2 infection model

**DOI:** 10.1101/2021.04.16.440101

**Authors:** Antonia Sophia Peter, Edith Roth, Sebastian R. Schulz, Kirsten Fraedrich, Tobit Steinmetz, Dominik Damm, Manuela Hauke, Elie Richel, Sandra Mueller-Schmucker, Katharina Habenicht, Valentina Eberlein, Leila Issmail, Nadja Uhlig, Simon Dolles, Eva Grüner, David Peterhoff, Sandra Ciesek, Markus Hoffmann, Stefan Pöhlmann, Paul F. McKay, Robin J. Shattock, Roman Wölfel, Ralf Wagner, Jutta Eichler, Wolfgang Schuh, Frank Neipel, Armin Ensser, Dirk Mielenz, Matthias Tenbusch, Thomas H. Winkler, Thomas Grunwald, Klaus Überla, Hans-Martin Jäck

**Affiliations:** Institute of Clinical and Molecular Virology, University Hospital Erlangen, Friedrich-Alexander University Erlangen-Nuremberg, Erlangen, Germany; Division of Molecular Immunology, Internal Medicine III, Nikolaus-Fiebiger-Center of Molecular Medicine, Friedrich-Alexander University Erlangen-Nuremberg, Erlangen, Germany; Division of Genetics, Department Biology, Nikolaus-Fiebiger-Center of Molecular Medicine, Friedrich-Alexander University Erlangen-Nuremberg, Erlangen, Germany; Department of Immunology, Fraunhofer Institute for Cell Therapy and Immunology IZI, Leipzig, Germany; Department of Chemistry & Pharmacy, Friedrich-Alexander University Erlangen-Nuremberg, Erlangen, Germany; Institute of Medical Microbiology and Hygiene, Molecular Microbiology (Virology), University of Regensburg, Regensburg, Germany; Institute of Medical Virology, University Hospital Frankfurt, Goethe University Frankfurt, Frankfurt, Germany; German Centre for Infection Research, External Partner Site, Frankfurt, Germany; Fraunhofer Institute for Molecular Biology and Applied Ecology (IME), Branch Translational Medicine and Pharmacology, Frankfurt, Germany; Infection Biology Unit, German Primate Center, Göttingen, Germany; Faculty of Biology and Psychology, Georg-August-University Göttingen, Göttingen, Germany; Department of Infectious Diseases, Imperial College London, Norfolk Place, London, UK; Bundeswehr Institute of Microbiology, Munich, Germany; German Center for Infection Research, Partner Site Munich

## Abstract

TRIANNI mice carry an entire set of human immunoglobulin V region gene segments and are a powerful tool to rapidly generate human monoclonal antibodies. After immunizing these mice against the spike protein of SARS-CoV-2, we identified 29 hybridoma antibodies that reacted with the SARS-CoV-2 spike protein. Nine antibodies neutralized SARS-CoV-2 infection at IC50 values in the subnanomolar range. ELISA-binding studies and DNA sequence analyses revealed one cluster of clonally related neutralizing antibodies that target the receptor-binding domain and compete with the cellular receptor hACE2. A second cluster of neutralizing antibodies binds to the N-terminal domain of the spike protein without competing with the binding of hACE2 or cluster 1 antibodies. SARS-CoV-2 mutants selected for resistance to an antibody from one cluster are still neutralized by an antibody from the other cluster. Antibodies from both clusters markedly reduced viral spread in mice transgenic for human ACE2 and protected the animals from SARS-CoV-2 induced weight loss. Thus, we report two clusters of potent non-competing SARS-CoV-2 neutralizing antibodies providing potential candidates for therapy and prophylaxis of COVID-19. The study further supports the use of transgenic animals with human immunoglobulin gene repertoires in pandemic preparedness initiatives.

## INTRODUCTION

Since December 2019 [1], SARS-CoV-2 has rapidly spread throughout the world, leading to more than 133,971,287 confirmed cases of COVID-19 and 2.902.493 million deaths [2]. Several vaccines have been developed and licensed in worldwide efforts at an unprecedented speed [3, 4]. In countries or age groups with high vaccine coverage, case counts have decreased substantially [5, 6]. For all licensed vaccines, the vaccine antigen is the ectodomain of the spike protein. Its receptor-binding domain (RBD) interacting with the cellular receptor human Angiotensin-converting enzyme 2 (hACE2) as well as its N-terminal domain (NTD) were identified as primary targets of the neutralizing activity of convalescent sera and monoclonal antibodies [7–19]. Clinical development of some of these monoclonal antibodies is ongoing, already providing evidence of their utility in therapeutic trials [20–22]. Some have since been licensed for the treatment of COVID-19 (reviewed in [23]).

After crossing species barriers, zoonotic viruses adapt to new species. Early after the new entry, adaptation to efficient transmission is most likely the most substantial selective pressure, explaining the rapid emergence of SARS-CoV-2 variants, such as those carrying the D614G mutation [24] or more complex mutational signatures in the spike protein [25, 26]. The higher the percentage of a population is that has overcome the first infection, the more robust the selective advantage of variants escaping the immune responses raised against the initial founder virus will be. Evidence that immune escape may already contribute to the emergence of new variants has been reported for the South African B.1.351 and the Brazilian P.1 and P.2 variants [27–29]. Since the antibody response to current mRNA vaccines precisely mimics the antibody response to natural infection [30], the vaccines’ efficacy against these variants may be impaired [27, 30–32]. Similarly, the neutralizing potency of monoclonal antibodies may be reduced or completely lost [15, 33, 34].

Concerning the loss of vaccine efficacy against emerging variants, it has been argued that mRNA and viral vector vaccines can be rapidly adapted to include the spike proteins of the variants of concern. While this can be expected to work for individuals with no prior exposure to SARS-CoV-2 or the COVID-19 vaccines, the outcome is less predictable in the presence of pre-existing antibody responses to the founder spike protein. As variants only differ in a few epitopes, immunizations with spike proteins of emerging variants in individuals previously exposed to the founder spike protein may lead to preferential boosting of pre-existing non-neutralizing antibodies to epitopes conserved between the two spike proteins. More targeted vaccine approaches employing structure-based designs may be needed but require reliable animal models for validation and optimization. Transgenic mice in which the coding sequences for variable regions of the immunoglobulin (Ig) murine heavy (H) and light (L) chains were replaced by coding sequences of variable regions of human H and L chains could be such a model.

Here we report the rapid isolation of fully human SARS-CoV-2 neutralizing monoclonal antibodies from immunized TRIANNI mice harboring the complete human antibody repertoire (patent US 2013/0219535 A1; Fig 1A). The neutralizing antibodies target the RBD or the NTD of the spike protein, providing a pair of non-competing human monoclonal antibodies for clinical development. Further characterization of the antibodies’ human VH gene usage and divergence from germline ancestor sequences supports the concept of mouse models with a human antibody repertoire in pandemic preparedness efforts and suggests the use of these models in structure-based optimization of vaccine antigens against emerging SARS-CoV-2 variants.

**Figure 1:**
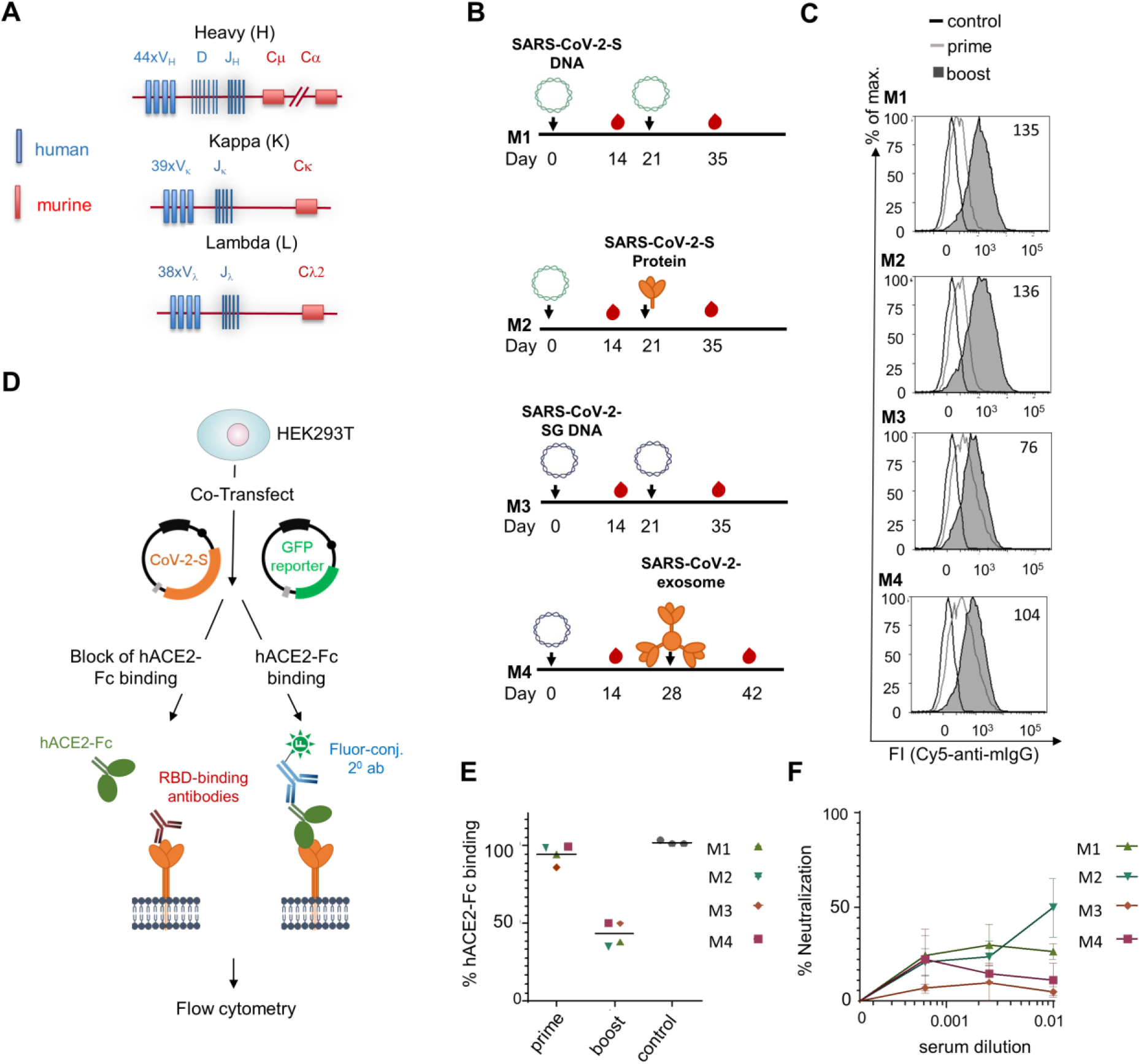
Immunization of TRIANNI mice for induction of SARS-CoV-2 neutralizing antibodies. TRIANNI mice harboring the entire human Ig variable region repertoire **(A)** were primed by intramuscular electroporation with expression plasmids for wild type SARS-CoV-2-S (M1, M2) or a hybrid SARS-CoV-2-S containing the intracytoplasmic domain of VSV-G (M3, M4) **(B)** Mice were boosted with the expression plasmids used for priming (M1, M3), soluble trimeric S protein (M2), or exosomes carrying the hybrid SARS-CoV-2-S protein (M4). **(C)** A flow cytometric assay assessed the binding of sera at a 1:200 dilution to the SARS-CoV-2-S protein with HEK-293T cells transiently expressing the S protein. Numbers indicate the relative mean fluorescence intensities of sera drawn two weeks after the booster immunizations. **(D)** Scheme of hACE2-Fc competition assay. HEK-293T cells expressing the SARS-CoV-2-S protein were incubated with hACE2-Fc fusion protein in the presence or absence of sera from immunized mice before staining with an AF647-labelled anti-human Fc antibody. **(E)** Competitive inhibition of hACE2-Fc binding to trimeric S protein by sera (1:200) from control mice and mice at the indicated time points after the first immunization. The mean percentage of binding as compared to control binding is shown (two experiments each performed in triplicates). **(F)** For the neutralization assay, Vero-E6 cells were infected with the SARS-CoV-2 isolate MUC-IMB-1 in the presence or absence of week 5 sera. SARS-CoV-2 infection was quantitated after 20 to 24 hours by staining with purified IgG from a convalescent COVID-19 patient and a fluorescence-labeled anti-human IgG using an ELISPOT reader. The mean and SEM of triplicates of one experiment are shown.

## RESULTS

### Immunization

Based on the published nucleotide sequence of SARS-CoV-2 [35], the first immunogens available early during the current pandemic were expression plasmids encoding the spike protein, the primary target of neutralizing antibodies to coronaviruses [36, 37]. As DNA vaccines also induce good T cell responses, we selected DNA vaccines as our priming immunogens. In addition to a DNA vaccine encoding the wild type S protein (SARS-CoV-2-S DNA), we generated a DNA vaccine in which the coding region of the cytoplasmic domain of SARS-CoV-2 was replaced by the cytoplasmic domain of the G protein of VSV (SARS-CoV-2-SG DNA). For SARS-CoV, it was shown that such a modification increases S protein expression at the cell surface and induces higher neutralizing antibody responses after immunization with exosomal vaccines [38]. The time interval between the DNA priming immunization and the booster immunization allowed for the production and purification of a recombinant S protein stabilized in the pre-fusion state (SARS-CoV-2-S protein) and an exosomal vaccine presenting a membrane-anchored form of the SARS-CoV-2-S protein (SARS-CoV-2-S Exo) (Fig. 1B).

TRIANNI mice were selected for these immunization experiments (Fig. 1A) because they produce antibodies with a human Ig variable (V) region and murine constant (C) regions. One unique feature of the TRIANNI mice is that the intronic regions between the exons of the variable segments of Ig genes are essentially of murine origin. This design aims to maintain proper regulation of expression and splicing of the V regions of the immunoglobulin exons as a prerequisite for proper affinity maturation. After selection of antibodies of interest, the human V region gene segments can be recombinantly fused to human C region gene segments allowing expression of fully human antibodies for clinical applications.

On February 14, 2020, groups of two TRIANNI mice were either immunized by intramuscular electroporation with the SARS-CoV-2-S DNA and or the SARS-CoV-2-SG-DNA (Fig. 1B). Three weeks later, the SARS-CoV-2-S DNA primed mice were either boosted with the SARS-CoV-2-S DNA or the SARS-CoV-2-S protein adjuvanted with MPLA liposomes. The SARS-CoV-2-SG DNA primed mice were boosted with SARS-CoV-2-SG or SARS-CoV-2 exosomes at weeks 3 and 4, respectively (Fig. 1B). Using a flow cytometric assay with cells transiently transfected with SARS-CoV-2-S DNA [38], antibody responses were analyzed two weeks after the priming and the boosting immunization (Fig. 1C). Antibodies to SARS-CoV-2-S were detectable in all four animals after the DNA priming immunization and further increased after the booster immunizations.

Sera of the immunized mice were also analyzed for competition with the binding of the cellular receptor hACE2 to the SARS-CoV-2-S protein (Fig. 1D, E). Two weeks after the booster immunization, all four sera reduced hACE2 binding. Priming with SARS-CoV-2-S DNA seemed to result in slightly higher competition activity. Sera from the mice primed with SARS-CoV-2-S DNA and boosted with SARS-CoV-2-S protein or SARS-CoV-2-S DNA also neutralized wild type SARS-CoV-2 at a 1:100 dilution by 50% and 27%, respectively (Fig 1F).

### Hybridoma screening for spike binding, ACE2 competing and neutralizing antibodies

The TRIANNI mouse (M2) with the best neutralizing activity was boosted with the adjuvanted SARS-CoV-2-S protein on week 6 and sacrificed five days later to generate hybridoma lines. A 1:100 dilution of the serum from this time point neutralized wild-type SARS-CoV-2 by 86%. Cells from the spleen and draining lymph nodes were fused with Sp2/0-Ag14 cells, and hybridoma cells were grown in 106 96-well plates seeded at a density of 1,000 cells per well. Between ten to 20 days after the fusion, antibodies secreted by the hybridoma cells were screened by flow cytometry for binding to the S protein. A total of 29 hybridomas could be identified that secreted antibodies binding to the S protein. These antibodies were designated TRES antibodies (<u>TR</u>IANNI-<u>E</u>rlangen anti SARS-CoV-2-<u>S</u>pike). The screening results of ten representative hybridoma supernatants containing TRES antibodies are displayed in Fig. 2A (for a list of all binding antibodies and their characteristics, see Table 1). Intracellular staining revealed that most hybridomas expressed the IgG2c subtype.

**Figure 2:**
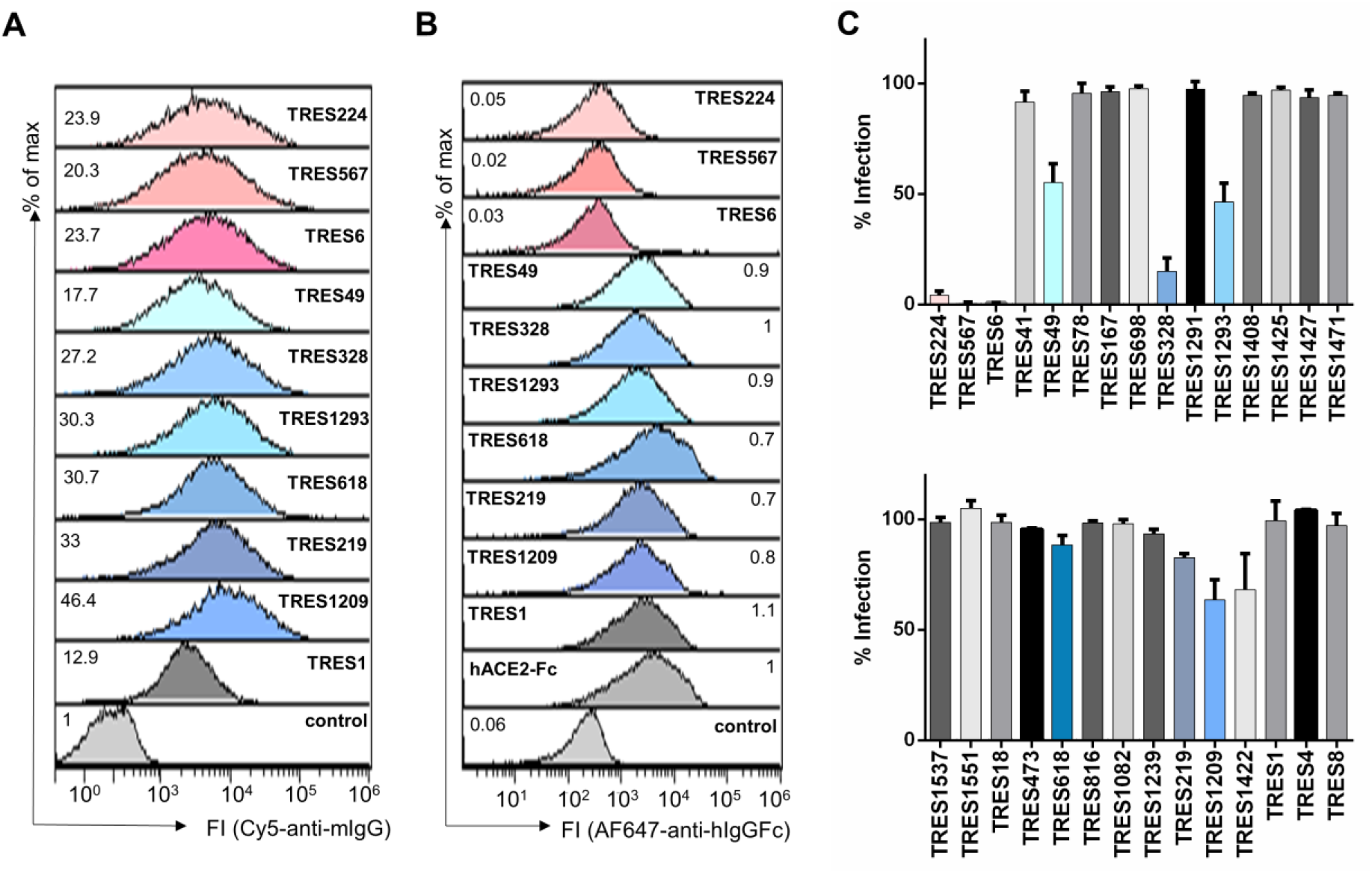
Screening of hybridoma supernatants. **(A)** The binding of antibodies from undiluted TRES hybridoma supernatants to HEK-293T cells expressing the SARS-CoV-2 spike protein was detected with a fluorescence-conjugated murine pan IgG antibody. Numbers indicate the relative mean fluorescence intensity. **(B)** Detection of hACE2-competing TRES antibodies as described in Fig. 1 D. Numbers depict the relative mean fluorescence intensity. **(C)** Vero-E6 cells were infected with the SARS-CoV-2 MUC-IMB-1 isolate in the presence or absence of undiluted TRES hybridoma supernatants. SARS-CoV-2 infection was quantified after 20 to 24 hours by staining as described in Fig. 1F. The mean and standard deviation of triplicates of one experiment are shown.

**Table 1:**
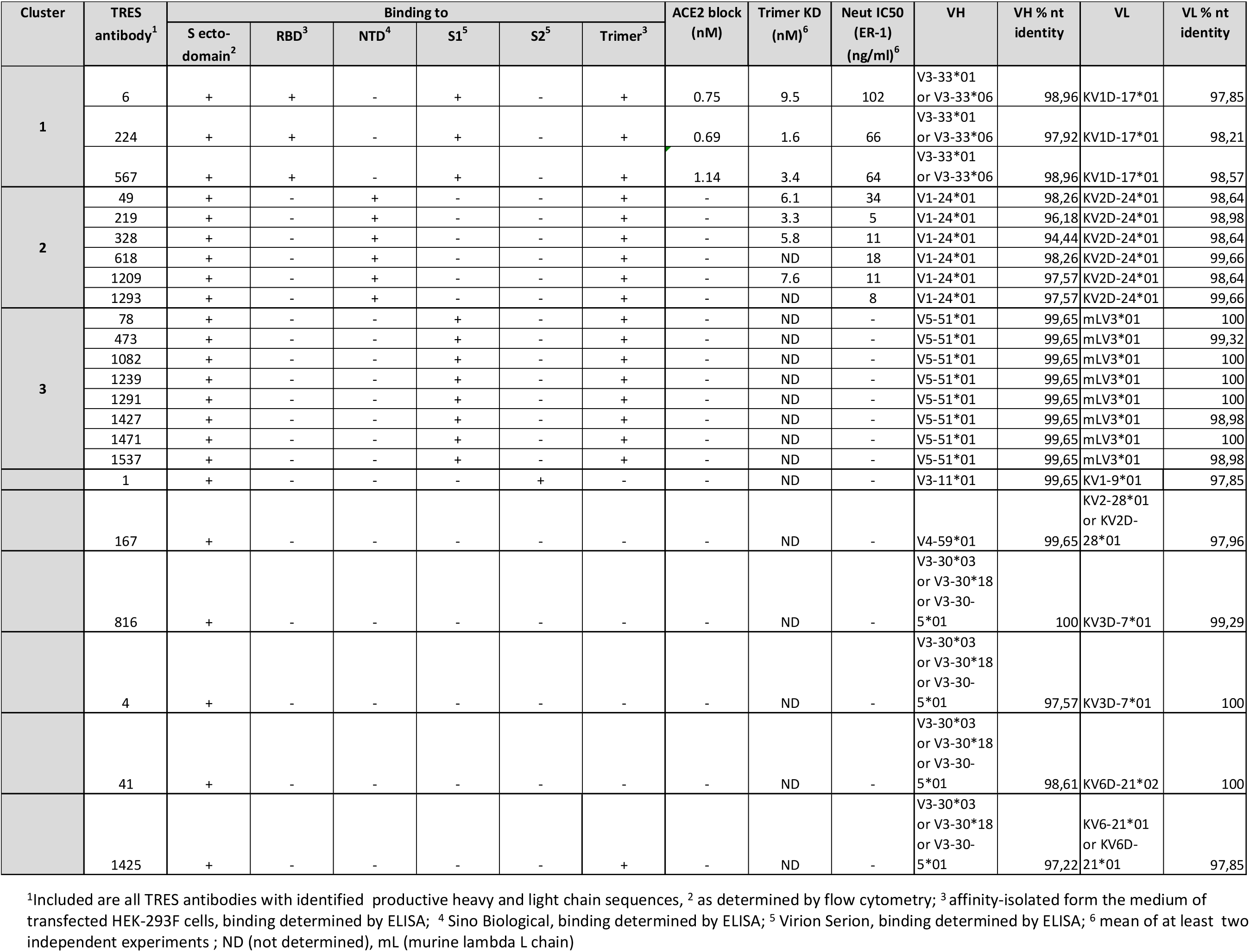
Overview: <u>TR</u>ianni-<u>E</u>rlangen anti-CoV2-<u>S</u>pike (TRES) antibodies

The binding antibodies were further characterized in an hACE2 competition assay. Three (TRES6, 224, 567) of the 29 binding antibodies were able to outcompete hACE2 binding very efficiently (Fig. 2B), indicating that these antibodies might bind the RBD and possibly neutralize SARS-CoV-2.

Subsequently, all supernatants were tested in a virus neutralization assay with SARS-CoV-2 MUC-IMB-1, an early isolate from the Webasto cluster in Bavaria [39, 40]. Supernatants from the hybridomas competing with ACE2-binding (TRES antibodies 224, 567 and 6) strongly neutralized SARS-CoV-2 (Fig. 2C) and indeed, an RBD-specific ELISA confirmed the RBD-binding of these three antibodies (Fig. 3A). Three additional hybridoma supernatants could be identified to inhibit virus infection by at least 40% (TRES49, 328, 1293), although they do not compete with hACE2 for binding to the SARS-CoV-2-S protein. They also do not bind to the RBD, recombinant S1 proteins or the S2 protein (Fig. 3A-C). TRES49, 328, 1293 bind to a stabilized S protein trimer and recombinant NTD (Fig. 3D, E). A similar binding profile was also observed for the TRES antibodies 618, 219 and 1209. Thus, these six antibodies differ in their binding profile from TRES6, TRES224 and TRES567. We also identified one hybridoma supernatant (TRES1) that strongly binds to S2 (Fig. 3B).

**Figure 3:**
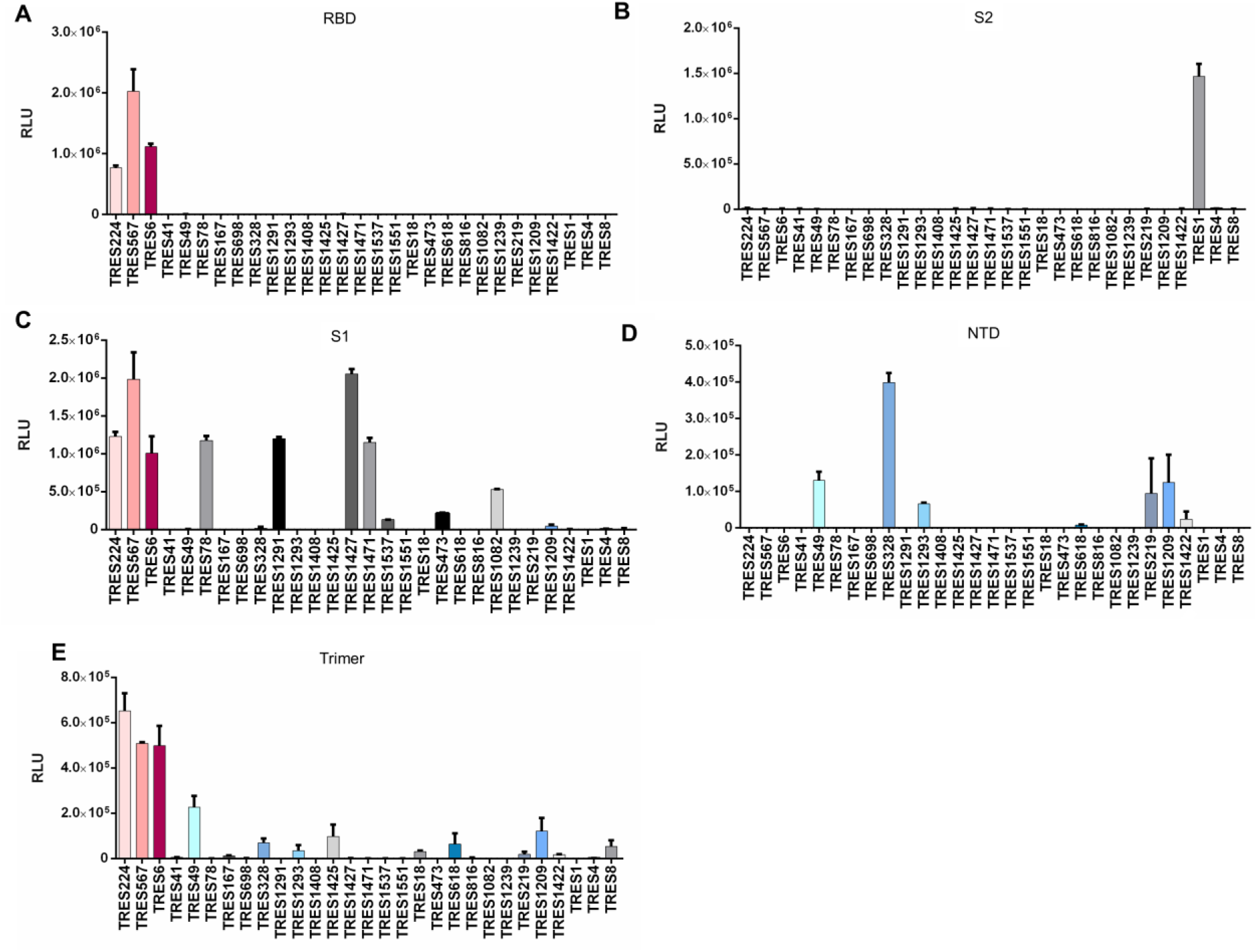
Binding pattern of hybridoma supernatants and purified antibodies. ELISA plates were coated with recombinant RBD (**A**), S2 (**B**), S1 (**C**), NTD (**D**), or SARS-CoV-2 spike protein stabilized in a prefusion state affinity-purified from medium of transfected HEK-293F cells (**E**) and incubated with TRES hybridoma culture medium. Bound antibodies were detected with HRP-labeled anti-mouse IgG antibodies and the RLUs measured. Mean and standard deviations of triplicates of one experiment are shown.

### Characterization of monoclonal TRES antibodies

Based on the screening results, hybridomas were subcloned, and antibodies from the supernatants of hybridoma subclones of the TRES antibodies 6, 224, 567, 49, 219, 328, 618, 1209, 1293 and 1 were purified and further characterized. Since the TRES antibodies 6, 224, and 567 compete with hACE2 for binding to RBD (Fig. 2A), the 50% effective concentration (EC50) was determined in an hACE2 competition ELISA t0 range from 0.75 to 1 nM (Fig. 4 A). Surface plasmon resonance (SPR) analyses further revealed K_D_ Values of purified RBD and NTD binding antibodies against the S protein stabilized in the prefusion state from 9.5 nM to 1.6 nM (Table 1).

**Figure 4:**
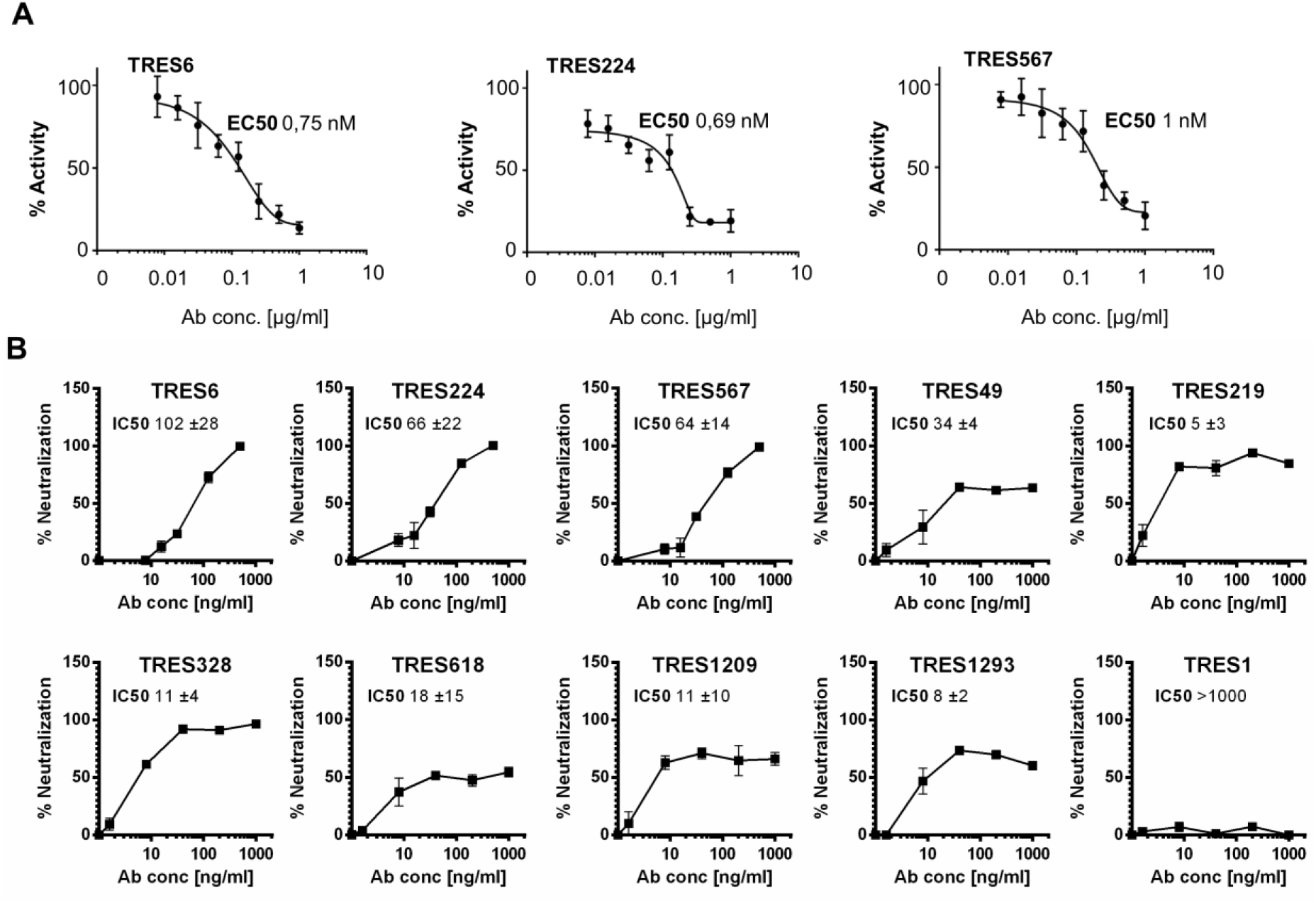
Characterization of monoclonal TRES antibodies. **(A)** ELISA-based hACE2-competition assay with TRES antibodies. Plates were coated with RBD and incubated with serial dilutions of TRES antibodies and soluble hACE2 (400 ng/ml). Bound hACE2 was quantitated with HRP-coupled antibodies against the hFcγ1-Tag of hACE2. Mean and standard deviation of quadruplicates of one representative experiment out of two are shown in the graphs. The mean EC50s of all experiments are also shown. **(B)** Neutralization assay. Vero-E6 cells were incubated with the CoV-2 ER-1 isolate with increasing concentrations of the respective TRES antibodies. SARS-CoV-2 infection after 20 to 24 hours was quantitated as described in Fig. 1 F. Graphs show the mean and SEM of triplicates of one representative experiment of at least three experiments. The mean IC50 and standard deviation, in ng/ml, of all experiments, is also given. IC50s were calculated with inhibitor vs. variable slope fitting curve with GraphPad Prism 7.02.

The neutralizing activity of purified antibodies was determined against a second wild type SARS-CoV-2 isolate, CoV-2-ER1 (Fig. 4B). In contrast to the MUC-IMB-1 isolate, CoV-2-ER1 contains the D614G mutation dominating the SARS-CoV-2 populations in Europe and the US [41, 42]. Antibodies that compete with hACE2 (TRES6, 224, and 567) had IC50s ranging from 64-102 ng/ml, while the IC50s of TRES49, 328, 618 and 1293 were lower, being between 5 and 34 ng/ml. Although the initial screening of hybridoma supernatants had not detected neutralization by TRES618, 219, 1209, the antibodies had an IC50 of 18 ng/ml, 5 ng/ml and 11 ng/ml, respectively. The S2-binding antibody TRES1 did not display neutralizing activity (Fig. 4B). When the antibodies were assessed in a cell-cell fusion assay, in which Vero-E6 and HEK-293T cells expressing the S protein were allowed to fuse, the RBD-binding antibodies seemed to inhibit cell-cell fusion more efficiently than the neutralizing antibodies not binding to RBD (Fig. S1). In general, cell-cell fusion inhibition required higher antibody concentrations and total inhibition of cell-cell fusion was not achieved (Fig. S1).

### Sequence analysis of variable regions of TRES antibodies

Sequencing of productive immunoglobulin variable regions expressed by hybridomas encoding TRES antibodies revealed a broad usage of V regions of H and L chains with a preference for κL chains (Table 1). Compared to the corresponding germline sequences, TRES VH and VL nucleotide sequences showed similarities as low as 94.44% and 97.85%, respectively, indicating the presence of somatic mutations and suggesting affinity maturation. At the same time, clonally related hybridomas could be identified. Consistent with their similar binding profiles (Fig. 3, 4), TRES6, 224, and 567 H and L chain exons contain the same V_H_DJ_H_ and V_L_J_L_ recombination joining sequences and are therefore designated cluster 1 antibodies. All three antibodies express the VH3-33*01 or VH3-33*06 and KV1D-17*01 gene segments (Table 1). An alignment of amino acid sequences of VH and VL of cluster 1 antibodies with the inferred germ line encoded antibody revealed an identical L chain with 5 non-silent somatic mutations in the antigen bindings loop (complementary determining regions, CDR) and one in the framework (FR) (Fig. S2) TRES6 and 567 utilize the same VDJ H chain exon with four somatic mutations in the framework. TRES224 contains two additional mutations in the CDR1 and CDR2 and an unusual change of a tyrosine that is conserved in 98% of all human VH segments. A second cluster comprises six TRES antibodies (Table 1; TRES49, 219, 328, 618, 1209 and 1293) that share the VHDJH and VLJL recombination joining sequences. Various somatic mutations are also present in the CDRs as well as in the FR of their VH and VL segments (Fig S2). The non-neutralizing antibodies of a third cluster bind to S1, but not to RBD, and are composed of an H chain with a human VH and a L chain with an unusual mouse VλX region (Table 1) [43], which could be explained by the fact that the mouse VλX region is still present in TRIANNI mice.

### Generation of fully humanized TRES antibodies

Gene segments encoding the human V exons of the H and L chains of all neutralizing TRES antibodies were fused with the human γ1 H and the κ L chain constant regions, respectively. Recombinant antibodies entirely composed of human Ig regions were produced in HEK-293F cells and purified by affinity chromatography. As expected, all recombinant TRES antibodies bound to the S protein in the flow cytometric assay (Fig. 5A).

**Figure 5:**
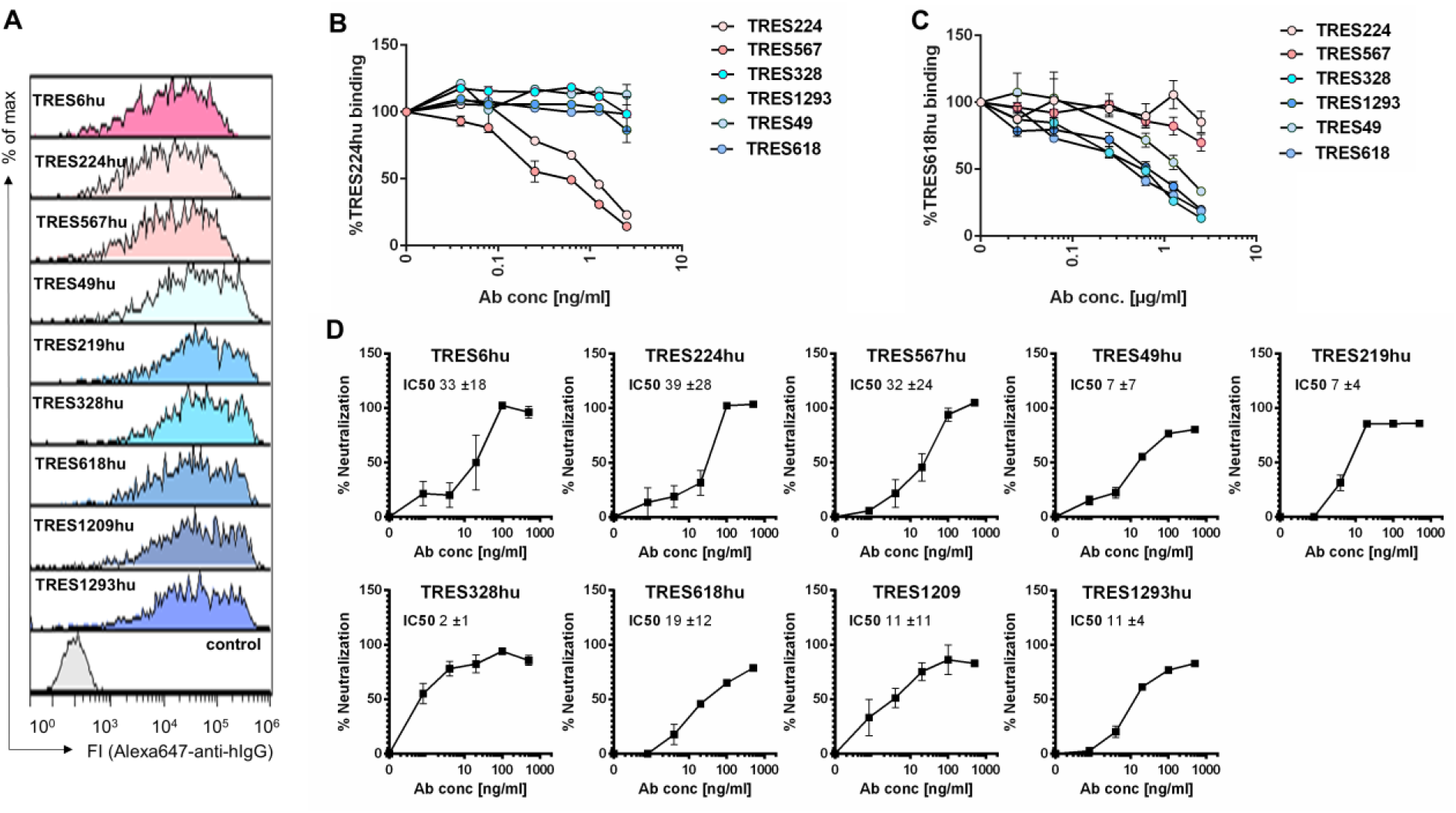
Characterization of recombinant human TRES antibodies. **(A)** Flow cytometric analysis of HEK-293T cells expressing the SARS-CoV-2-S protein and stained with recombinant humanized IgG1 TRES (TREShu) antibodies and a fluorochrome-labeled secondary antibody against human IgG-Fc. A non-S binding human antibody served as a negative control. **(B, C)** HEK-293T cells expressing the SARS-CoV-2 spike protein were incubated with recombinant TRES antibodies with a human Fcγ1 region and serially diluted TRES hybridoma antibodies with a murine Fcγ. Bound recombinant human TRES224hu (B) or TRES618hu (C) were detected with a mouse Alexa647-labeled antibody directed against the human Fcγ region. The mean percentages of binding and SEM of one experiment performed in triplicates are shown. **(D)** The SARS-CoV-2 neutralizing activity of the human recombinant TRES antibodies was analyzed as described in Fig. 4B. Shown are means and SEM of triplicates of one representative experiment out of three. Also given are the mean and standard deviation of IC50s, given in ng/ml, of the three independent experiments, calculated as described in Fig. 4B.

To explore a potential overlap of epitopes between TRES antibodies of cluster 1 and cluster 2, cells expressing the SARS-CoV-2-S protein were incubated with TRES224hu or TRES618hu in the presence of increasing concentrations of TRES224, 567, 49, 328, 618 and 1293 hybridoma antibodies containing mouse C regions (Fig. 5B, C). No competition between the cluster 1 and cluster 2 antibodies could be observed. In contrast and as expected, binding of the recombinant TRES224 with human C regions (TRES224hu) was efficiently blocked by TRES567 and 224 (Fig. 5B), indicating that these two neutralizing antibodies from cluster 1 recognize the same epitope. Accordingly, the binding of TRES618hu was blocked by TRES49, 328, 618 and 1293, verifying that cluster 2 antibodies bind the same epitope. Most importantly, however, the humanized TRES antibodies also neutralized SARS-CoV-2 with similar IC50s as the parental hybridoma TRES antibodies (Fig. 5D), confirming that the identified antibody sequences confer neutralization.

### Breadth of neutralization and escape mutants

Neutralization assays (Fig. 6A) and spike protein binding assays (Fig. S3) were also performed for two emerging SARS-CoV-2 variants of concern and three antibodies from each cluster. The IC50s of the cluster 1 antibodies against the B.1.1.7 variant were in the range of 6 to 49 ng/ml (Fig. 6A, left panel) and were similar to those for the wild-type virus (Fig. 5B). IC50s consistently below 10 ng/ml (Fig. 6A, right panel) indicate that the B.1.351 isolate might be more susceptible to neutralization by cluster 1 antibodies than the wild-type virus (see Fig. 5D). In contrast, cluster 2 antibodies lost their neutralizing activity against both variants of concern (Fig. 6A).

**Figure 6:**
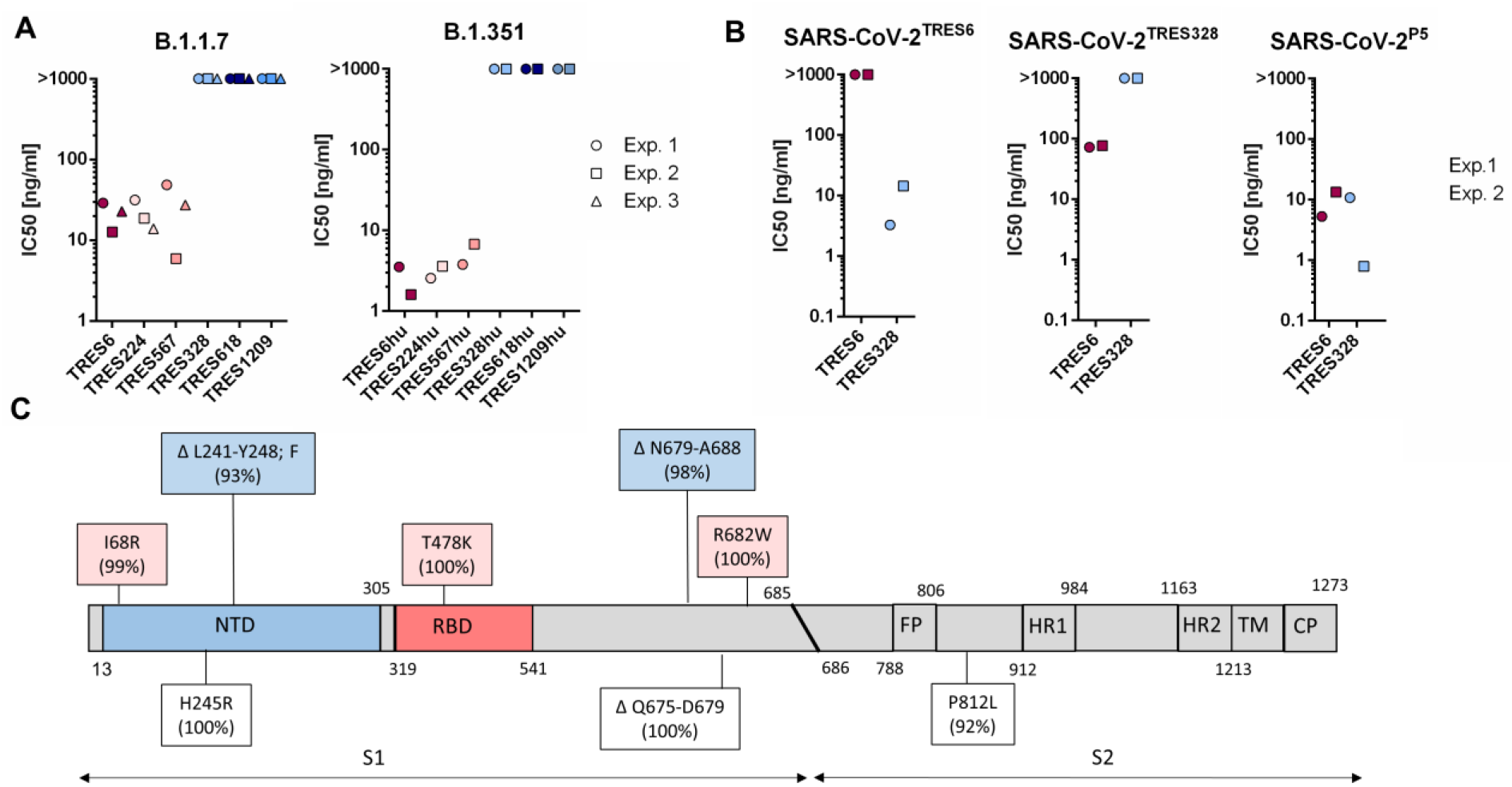
Breadth of neutralization by TRES antibodies and emergence of escape mutants. **A)** Neutralizing activity of hybridoma (TRES) or recombinant human TRES (TREShu) antibodies towards B.1.1.7 or B.1.351 determined as described in Fig. 4B. Shown are IC50s calculated from two to three experiments each performed in triplicates. **(B)** Emergence of escape mutants in cell culture during 5 passages on Vero-E6 cells in the presence of TRES6 or TRES328 in the absence or presence of increasing antibody concentrations. IC50s were determined in two independent experiments performed in triplicates as described previously in Fig. 4B. **(C)** Mutations in the S-Protein (not scaled) of the SARS-CoV-2 TRES6 (pink) and TRES328 (blue) escape mutants and the P5 variant passaged without antibody (white) as determined by whole-genome sequencing. The percentage describes the frequency of the mutations in the virus population. Exp (Experiment), NTD (N-terminal domain), RBD (receptor-binding domain), FP (fusion peptide), HR (heptat repeat), TM (transmembrane domain), CT (cytoplasmic tail).

To further explore the possibility for de novo emergence of resistant variants to cluster 1 and cluster 2 antibodies, the CoV-2 ER-1 isolate carrying the D614G mutation was passaged in increasing concentrations of TRES6 and TRES328. After 5 passages, neutralization assays were performed. While the passaged virus became resistant to antibody neutralization when the antibody was present during passaging, the virus remained sensitive to the TRES antibody when the antibodies were not included during passaging (Fig. 6B). Whole-genome sequencing of the passaged SARS-CoV-2 viruses revealed the emergence of a TRES6 antibody escape variant with an I68R mutation in the NTD and a T478K mutation in the RBD (Fig. 6C, displayed in pink). In contrast, the TRES328 escape variant harbored a L241-Y248 deletion with a phenylalanine insertion in the NTD (Fig. 6C, displayed in blue). Additional point mutations and deletions were observed in proximity to the S1-S2 cleavage sites. These mutations probably reflect adaptation to the replication in the Vero-E6 host cells [44] since these sites were also deleted in the virus variant (P5) passaged in the absence of a neutralizing antibody (Fig. 6C, displayed in white). Additional mutations were detected in the P5 variants at the positions H245 and P812 probably also reflecting viral adaptation to cell culture.

### Efficacy in a murine challenge model

To assess whether the antibodies generated in this study can confer protection from disease, we evaluated the efficacy of one antibody from each cluster in a stringent hACE2 transgenic mouse [45] viral challenge model under post-exposure prophylactic settings. Three groups of mice (n=12/group) were infected intranasally with 300 FFUs of SARS-CoV-2 (MUC-IMB-1). One day later, mice were injected intravenously with 5.25 mg/kg body weight of the cluster 1 antibody TRES6, the cluster 2 antibody TRES328, or the isotype-matched control antibody TRES480.

Six animals from each group were sacrificed on day 4 or 10 or according to humane endpoints, and viral loads were determined in the lung, BAL, brain, liver and spleen. Both antibodies reduced the amount of viral RNA in the lung and BAL samples approximately 30-to 200-fold after four days (Fig. 7A) or approximately 100-o 450-fold after 10 days (Fig. 7B). Viral load in the other organs was reduced close to the level of detection. Interestingly, the titer of infectious virus in the BAL samples from TRES6 and TRES325 treated animals was below the detection level and, therefore, at least 1000-fold lower than in mice receiving the control antibody (Fig. 7C). Compared to the isotype control, TRES480, treatment of mice with TRES6 and TRES328 prevented the loss of body weight induced by the viral infection (Fig. 7) and reduced clinical symptoms assessed by a clinical score (Fig. 7E). Importantly, none of the TRES antibody-treated mice reached clinical endpoints requiring euthanasia, while 4 of 6 SARS-CoV-2-infected mice receiving the isotype control antibody TRES480 had to be euthanized (Fig. 7F).

**Figure 7:**
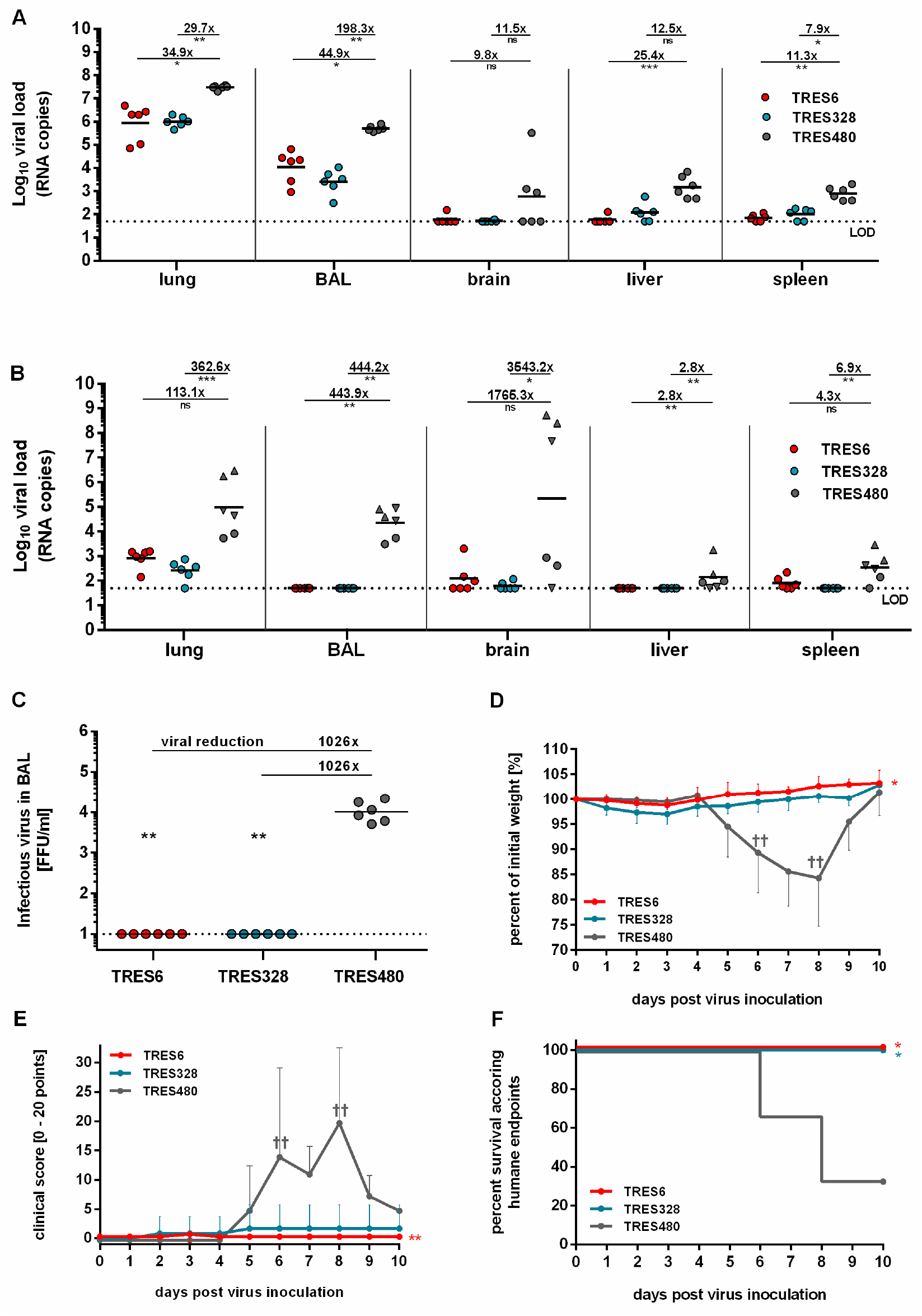
Efficacy of TRES6 and TRES328 in a post-exposure prophylactic model. Reduction of viral load in hACE2-transgenic mice treated with TRES6, TRES328 or TRES480 isotype control antibody. Mice were inoculated intranasally with 300 FFU of SARS-CoV-2 on day 0. One day later, mice were treated intravenously with 5.25 mg/kg TRES6, TRES328 or TRES480 control antibody. Viral loads were determined on day 4 **(A)** or day 10 **(B)** after virus inoculation by RT-qPCR in the indicated organ samples. Data points represent the viral copy number of individual animals with the geometric means of each group depicted as lines, circles (•) indicate the survival of 4 or 10 days post-infection, and triangles indicate euthanized mice according to humane endpoints at day 6 (▲) or day 8 (▼). Calculated reduction of viral RNA is shown in comparison to the TRES480 control group. **(C)** Infectious virus load in BAL samples from antibody and isotype treated mice. Infectious virus was measured by focus-forming assay. **(D)** Body weight and **(E)** clinical score of antibody treated and isotype treated mice. Animals reaching humane endpoints were euthanized and are marked by a cross (†). **(F)** Survival curve of antibody treated and isotype control treated animals. Percent survival as the fraction of animals surviving humane endpoints (Kaplan-Meier analysis). Statistical analysis of the presented data was performed by Kruskal-Wallistest (one way ANOVA) and Dunn’s Pairwise Multiple Comparison Procedures as post hoc test in comparison to the TRES-480 control (ns: non-significant, *: p < 0.05, **: p < 0.01, ***: p < 0.001).

## DISCUSSION

By immunizing TRIANNI mice with a DNA prime protein boost regimen, 29 monoclonal antibodies were obtained that bind to the S protein of SARS-CoV-2 expressed on the cell surface of transfected HEK-293T cells. The use of immunization and screening approaches based on the expression of the wildtype S protein avoids the need for prior knowledge of domains targeted by neutralizing antibodies and allows to obtain a panel of antibodies binding to different epitopes. Indeed, different binding patterns were observed, including antibodies that i) bind to the RBD and compete with ACE2 binding, ii) bind to the S1 subunit without binding to RBD, iii) only bind to the ectodomain of the S protein stabilized in a trimeric form and the NTD and iv) bind to the S2 subunit. In those mice that we first challenged with SARS-CoV-2 and subsequently treated with the monoclonal antibodies, we could also assess any potential disease-enhancing properties. This issue has been suggested as a potential outcome of infections occurring in the presence of vaccine induced antiviral antibodies not providing sterilizing immunity [46]. However, no evidence for such an enhancing effect could be observed. A combination therapy with an antibody from cluster 1 and cluster 2 seems especially attractive since this may extend the neutralization breadth and enhance the genetic barrier for the emergence of antibody-resistant SARS-CoV-2 variants. This assumption is further strengthened by the observation that the B.1.1.7 and the B.1.351 variants maintain neutralization sensitivity to at least one of the two clusters of antibodies. In addition, virus escape mutants from one antibody of each cluster are still sensitive to a representative TRES antibody from the other cluster.

Most of the neutralizing antibodies in convalescent COVID-19 patients are close to the germline sequence [47]. In contrast, the TRES antibodies had mutations in the VH and VL regions including the CDRs (Fig. S. 2) suggesting that affinity maturation can indeed occur in TRIANNI mice after a DNA prime protein boost immunization regimen. Although mice encoding a full [11] or partial [48] human immunoglobulin repertoire have been used previously to generate monoclonal antibodies targeting the spike protein of SARS-CoV-2, direct evidence for affinity maturation, in the context of SARS-CoV-2, has not yet been provided. Using a three-week immunization regimen consisting of a single intradermal priming immunization with DNA encoding the S protein and booster immunizations with an RBD-Fc fusion protein in VelocImmune™ mice, Hansen et al. showed induction of RBD-specific antibody responses and also reported the isolation of neutralizing RBD-binding antibodies from B cells that were sorted based on RBD binding [11]. Wang et al., performed sequential immunizations of H2L2™ (Harbour Antibodies) mice with spike ectodomains of different human coronaviruses and derived a monoclonal antibody that cross-neutralized wildtype SARS-CoV-2 moderately with an IC50 of 570 ng/ml [48].

Although extensive affinity maturation does not seem to be necessary for the formation of potent neutralizing antibodies against SARS-CoV-2 [47], this may be relevant for elicitation of neutralizing antibodies against other pathogens as exemplified by HIV-1 [49–51]. As observed for four of seven other potent neutralizing antibodies targeting the NTD and recovered from convalescent patients [7], our TRIANNI mouse-derived NTD antibodies also use the VH1-24 gene segment. Therefore, transgenic animal models supporting affinity maturation of antibodies with human variable regions should be an essential part of future pandemic preparedness efforts and be explored in structure-guided vaccine antigen design against SARS-CoV-2.

Besides the success in generating potent neutralizing antibodies that also confer protection in the hACE2 mouse model, a key criterion for applying TRIANNI mice in future pandemic responses is the time needed to obtain such antibodies. The immunization schedule we used took seven weeks until hybridomas were generated. Growing the hybridomas, their screening and subcloning took approximately six weeks. Sequence analyses from these hybridomas and gene synthesis for the generation of fully human antibodies took another six weeks. Accelerating the development time would certainly be desirable. Whether shortening the immunization schedule still results in potent neutralizing antibodies is difficult to predict and may depend on the degree of affinity maturation needed to generate such antibodies. However, instead of generating hybridomas, antigen-specific memory B cells or germinal center-selected surface IgG-positive plasmablast could be stained with fluorescence-labeled antigen and sorted by flow cytometry. Paired VH and VL sequences from single cells could then be amplified by PCR and directly cloned into expression plasmids, essentially as described for memory B cells from convalescent COVID-19 patients [10, 52]. Transiently expressed recombinant antibodies could then be screened for binding and neutralization and sequenced, probably shortening the development time by several weeks.

## MATERIAL AND METHODS

### Mice

The TRIANNI C57/Bl6 mouse line HHKKLL was established in cooperation with TRIANNI (Patent US 2013/0219535 A1). Mice were maintained under specific pathogen-free conditions in the Franz-Penzoldt Center animal facility of the University of Erlangen-Nürnberg following institutional and national regulations and the Federation of European Laboratory Animal Science Associations. All animal experiments were approved by the Regierung von Unterfranken (TVA 55.2.2-2532-2-961).

### Immunization of TRIANNI mice and the establishment of SARS-CoV-2-specific hybridoma clones

Two 9-12-week old TRIANNI mice were primed with pCG1-CoV-2019-S ([53] designated SARS-CoV2-S DNA) encoding wild type SARS-COV-2 spike protein (position 21580 – 25400 from Genbank NC_045512). Two additional TRIANNI mice were primed by SARS-CoV-2-SG DNA encoding a chimeric protein in which the intracytoplasmic domain of SARS-CoV-2-S was replaced by the intracytoplasmic domain of VSV-G as previously described for SARS-CoV [38]. The DNA vaccines were delivered by intramuscular electroporation as previously described [54]. Briefly, a total of 30 μg DNA diluted in 60 μl PBS were injected in both hind legs under constant isoflurane (CP-Pharma, Burgdorf, Germany) anesthesia. Immediately after the intramuscular injection, electrical impulses were applied to the injection site with a TriGrid electrode array (provided by Drew Hannaman, Ichor Medical Systems, Inc., San Diego, USA). Mice were boosted intramuscularly either with i) 5 μg of the S protein of SARS-CoV-2 stabilized in a pre-fusion conformation (designated SARS-CoV-2-S protein) and adjuvanted with 25 μg Monophosphoryl Lipid A (MPLA) liposomes (Polymun Scientific GmbH, Klosterneuburg, Austria) into the hind leg, ii) by electroporation of the DNA vaccines used for priming, or iii) with exosomes purified from HEK-293T cells transiently transfected with SARS-CoV-2 DNA as described previously [38]. Details for the immunization schedule are given in Figure 1B. Mice were bled by punctuation of the retro-orbital sinus using non-heparinized capillaries (Hirschmann Laborgeräte, Eberstadt, Germany). Serum was collected, inactivated for 30 min at 60°C and kept at −20°C for long-term storage.

One of the immunized mice received a second booster immunization with SARS-CoV-2-S protein at week 5. Five days later, spleen and inguinal lymph nodes were harvested and fused with the azaguanine-resistant SP2/0 Ag14 hybridoma line (ATCC#CRL-1581) using the PEG (Sigma Aldrich, Taufkirchen, Germany) method [55]. Briefly, 2×10^8^ spleen cells and inguinal lymph node cells (about 1×10^8^ B cells) were mixed with 10^8^ Sp2/0 cells and washed 3 times with RPMI1640 medium (Gibco, Thermo Fisher Scientific, Waltham, USA) without FCS. 2 ml PEG was added dropwise within one minute to the suspended cell pellet. The mix of fused cells was resuspended in R10+ medium (RPMI 1640 with 10% FCS, 0.05 mM mercaptoethanol, 2 mM L-glutamine, 1 mM sodium pyruvate, and 1% penicillin/streptomycin; (all from Gibco, Thermo Fisher Scientific, Waltham, USA); enriched with HAT (Gibco, Thermo Fisher Scientific, Waltham, USA) and OPI Media supplement (Sigma Aldrich, Taufkirchen, Germany). Cells were seeded into 96-well plates with approximately 1×10^4^ cells per well. 10-20 days post-fusion, hybridoma clones were tested for spike-binding antibodies. Positive clones were expanded in R10+ supplemented with HT (Sigma Aldrich, Taufkirchen, Germany) and subcloned by the limiting dilution method.

### Flow cytometric screening of hybridoma clones for SARS-CoV-2-specific antibodies and Ig isotypes

To detect antibodies that bind to SARS-CoV-2-S protein, HEK-293T cells were co-transfected with SARS-CoV-2-S DNA or pCG1-SARS-Fra1-S (CoV-1; position 21492 to 25259 Genbank AY291315SARS-CoV) and a GFP reporter plasmid (e.g., pEGFP-C1) using the PEI method as described previously [56]. Briefly, 2×10^7^ semi-confluent HEK-293T in 10 ml D1.5+ (DMEM+ 5% glutamine + Pen/Strep + 1.5% FCS) cells were incubated with 2.5 ml of a transfection mix in D0 (DMEM+ 5% glutamine + Pen/Strep) containing 30 μg of pCG1-CoV-2019-S or pCG1-SARS-Fra1-S and 10 μg of pEGFP-C1 plasmids and 180 μl of PEI (1mg/ml). After 6-8 hours at 37°C, the medium was changed to D10+ (DMEM+ 5% glutamine + Pen/Strep + 10% FCS). 48-72 hours post-transfection, cells were harvested without trypsin treatment, washed in FACS buffer (PBS with 0.5% bovine serum albumin and 1 nmol sodium azide) and used for binding assays or frozen in aliquots at −70°C. 10^5^ thawed or freshly transfected cells were incubated first in wells of a 96-well plate with 100μl undiluted hybridoma supernatant or 100 μl mouse serum (1:200 dilutions in R10+ medium), and bound antibodies were detected with a mix of Cy5-conjugated goat anti-pan-mouse IgG-Cy5 conjugated goat anti-mouse (Southern Biotechnology, Birmingham, USA, #SBA-1030-15) antibody. Cells were washed in FACS buffer and analyzed with an Attune Nxt, CytoFlex or a Gallios flow cytometer (Thermo Fisher Scientific, Waltham, USA or Beckman Coulter, Brea, USA, respectively) and evaluated with Flow Logic™ (IInivai Technologies Mentone, Australia).

Hybridoma cells were stained intracellularly for the assessment of the monoclonality of the colonies and the determination of the IgG subtype. Briefly, 10^5^ cells were fixed for 20 min in 2% paraformaldehyde (Morphisto, Frankfurt am Main, Germany) diluted in PBS, washed twice with FACS buffer, resuspended in permeabilization buffer (0.5% Saponin Sigma Aldrich, Taufkirchen, Germany, in FACS buffer) containing fluorochrome-conjugated murine H and L isotype-specific antibodies (antimouse IgG1-APC # 550874 and anti-mouse IgG3-bio # 020620 from BD, Franklin Lakes, USA, antimouse IgG2b-PE SBA-1090-09 and anti-mouse IgG2c-bio SBA-1079-08 from Southern Biotech, Birmingham, USA, anti-mouse Ig light chain lambda APC # 407306 and anti-mouse IgG light chain kappa PE both from Biolegend, San Diego, USA) and incubated for 1 hour at 4°C.

### FACS and ELISA based ACE2 competition assays

For quantitative flow-based hACE2 competition assays, 10^5^ SARS-CoV-2 spike-transfected HEK-293T cells were incubated with 50 μl of hACE2-Fc (250 ng/ml) composed of the ectodomain of human ACE2 fused in frame to the Fc region of human IgG1 and purified as described previously [56]. Subsequently, 50 μl undiluted hybridoma supernatant or mouse sera, diluted 1:200, were added and the cells were incubated on ice for an additional 30 minutes. Cells were washed in FACS buffer and stained on ice for 30 minutes with an Alexa647-labelled anti-human IgG-Fc antibody (Biolegend, San Diego, USA, #409320). Alexa647 mean fluorescence intensities were determined for GFP-positive HEK-293T cells with a FACS Attune Nxt and analyzed with the software Flow Logic™ (IInivai Technologies Mentone, Australia).

For ELISA-based hACE2 competition, 96-well microtiter plates were coated overnight at 4°C with 20ng/well (400 ng/ml) recombinant RBD that was purified from the culture medium of transfected HEK-293F cells. hACE2 and RBD proteins were generated by transient transfection of HEK-293F cells. For this the following plasmids were transfected: The plasmid pCEP4-hACE2ecto-Fc encodes a fusion protein containing the N-terminal ectodomain of hACE2 (AA 18-738 NCBI RefSeq: NP_001358344.1) the Hinge-CH2-CH3 region of human IgG1 and a C-terminal Myc-tag and 6xHis-tag. The vector encoding pSARS-CoV-2-RBD was constructed by inserting a commercially synthesized RBD fragment (AA 319-541, GenBank: QHD43416.1) with an N-terminal IgKappa signal sequence (METDTLLLWVLLLWVPGSTG) and C-terminal TwinStrep- and 3xFLAG-tags (GeneArt, ThermoFisher, Regensburg, Germany) into a pcDNA3.4(+) vector backbone. SARS-CoV-2 RBD and hACE2 proteins were purified from HEK-293F supernatants via affinity chromatography with Strep-Tactin^®^XT 4Flow agarose (IBA, Göttingen, Germany) following the manufacturer’s instructions. Eluted proteins were dialyzed against PBS in Slide A_Lyzer Dialysis Cassettes (Thermo Fisher Scientific, Waltham, USA) and stored in aliquots in 50% glycerol at −20°C. Protein purity was assessed by SDS PAGE followed by Western blot or Coomassie staining as described previously [57]. Protein concentrations were determined by OD at 260 nm and verified by a Bradford assay (Thermo Fisher Scientific, Waltham, USA). Plates were washed with PBS/0.05% Tween-20 and blocked with 275 μl per well of PBS with 2% BSA at room temperature (RT) for 1-3 hours. Competitional binding in the ELISA set up was achieved by applying 50 μl/well of 0.25 μg/ml human ACE2-Biotin (Acro Biosystems, Beijing, China, # A011-214) followed by 50 μl/well serially 2-fold pre-diluted TRES antibodies (2 μg/ml start concentration). This was incubated at RT for 1-2 hours. Wells were washed, 50 μl/well HRP-coupled Streptavidin (0.25 μg/ml, Merck Millipore, Darmstadt, #OR03L) were added and incubated at RT for 1-2 hours. After another washing step, HRP-bound Streptavidin was detected by adding 50 μl/well TMB substrate (BD Bioscience, Heidelberg, Germany, #555214). The reaction was stopped using 50 μl/well 0.5 M H_2_SO_4_ solution, and the OD at 450 nm was determined in a Spectra Max 190 (BMG Labtech, Ortenberg, Germany). EC50 values were calculated by plotting hACE2 activity against antibody concentrations and applying a 4-parameter curve fit using GraphPad Prism 7.02 (San Diego, USA).

### Detection of SARS-CoV-2 spike binding antibodies by ELISA

96-well microtiter plates were coated over night at 4°C with 100ng per well of recombinant NTD (Sino Biological; Beijing, China #40591-V49H), SARS-CoV-2 Spike S2 or S1 Glycoprotein (Virion\Serion, Würzburg, Germany), RBD or a S protein stabilized in a prefusion state (both affinity purified from the supernatant of HEK-293F cells, as described previously [58, 59]). The plates were washed and blocked with 5% skimmed milk for 1 hour at room temperature and then incubated with purified TRES antibodies or hybridoma supernatants for 1 hour. Next, the plates were washed, and goat-anti-mouse HRP (Dianova, Hamburg, Germany; #115-035-146) was added and incubated for 1hour. The plates were washed, and RLU (relative light units) were detected with the multilabel plate reader Victor X4 (Perkin Elmer, Hamburg, Germany).

### Determination of antibody affinity by SPR

Antibodies were captured on a Protein G Chip (Cytiva Lifesciences Protein G Chip; Marlborough, USA) to reach a response level of 500 RU. Following a kinetic titration was performed (t_Association_ = 180 s, t_Dissociation_ = 360 s) using a three-fold serial dilution of the S protein and spanning 5 concentrations. The chip surface was regenerated using 10 mM glycine, pH 1.5, as recommended by the manufacturer. The data was analyzed with Langmuir Kinetics and the BIAcore x100 Evaluation Software 2.1. The measurements were performed in triplicates on a Cytiva Lifesciences BIAcore X100 Plus. The S protein used was stabilized in a prefusion state as described [58]. It contained an N-terminal TPA signal peptide and a trimerization foldon followed by a His Tag at the C-terminus. The S protein was purified from transiently transfected HEK-293F cells as described previously [59].

### Cloning and sequencing of Ig V exon sequences

According to the manufacturer instructions, RNA was isolated from hybridoma cell lines with the RNAeasy Mini Kit (Qiagen, Aarhus, Denmark). 500 ng total RNA was used with the Template Switching RT Enzyme Mix (NEB Ipswich, USA) to generate 5’ RACE cDNA. Briefly, 1 μM of dT40 VN reverse primer and 1mM dNTP Mix were incubated for 5 minutes at 70°C together with 500 ng RNA in a 6μl reaction. The RT Enzyme mix and buffer were added to the reaction together with a Template Switch Oligo containing 3’ riboguanosines (rGrGrG; for primer sequences see Supplemental Table 1). The reverse transcription/template-switching reaction was incubated for 90 min at 42°C and inactivated at 85°C for 5 min. The template cDNA was then PCR amplified with 5’ primers annealing to the template switch oligo and 3’ primers specific for the respective Ig constant region of γ, κ and λ and for V_H_/V_K_/V_Λ_ with Q5 Hot Start polymerase (NEB, Ipswich, USA). PCR products were run on agarose gels, and 500-700 bp bands were excised and purified (QIAquick Gel Extraction Kit, Qiagen, Aarhus, Denmark). The purified PCR products were then sequenced with gene-specific primers or cloned with the CloneJET PCR Cloning Kit (Thermo Fisher Scientific, Waltham, USA) and introduced into competent DH5α E. coli cells. For each PCR product, at least four clones were sequenced to determine a consensus sequence. The consensus sequences were analyzed using VDJsolver [60] and IMGT/V-Quest [61].

### Construction of human IgG expression vectors

The cloning strategy was adapted from Tiller et al [52]. The vectors AbVec-hIgG1 [GenBank: LT615368.1] and AbVec-hIgKappa [GenBank: LT615369.1] were linearized by AgeI/SalI and AgeI/BsiWI digestion, respectively, separated by agarose gel electrophoresis and purified with the QIAquick Gel Extraction Kit (Qiagen, Aarhus, Denmark). Doublestranded DNA fragments (gBlocks) covering the respective mature V_H_ or V_L_ sequence, additional amino acids (VHS) to complete the leader sequence and 23bp overlap sequences at the 5’ and 3’ end were synthesized (IDT, San Jose, USA). The linearized vector and the synthesized fragment in a 1:2 molar ratio were assembled with the NEBuilder HiFi DNA Assembly Kit (NEB, Ipswich, USA) at 50°C for 30 min. Competent DH5α E. coli were transformed with 2 μl of the assembly mix and plated on selective agar plates. The plasmid sequences were confirmed by Sanger sequencing (Macrogen Europe, Amsterdam, The Netherlands).

### Purification of antibodies from culture supernatants

Murine antibodies were purified from serum-free cell supernatants from hybridoma via Protein-G affinity chromatography. Humanized antibodies were produced by transfecting HEK-293 F (Thermo Fisher Scientific, Waltham, USA) with the respective plasmids with the FreeStyle™ 293 Expression System according to the manufacturer’s instructions. 4 days post-transfection, proteins were affinity-purified. Hybridoma cells were grown in serum-free CD hybridoma medium (ThermoFisher Scientific, Waltham, USA) supplemented with 4 mM L-Glutamine to a cell density of about 1-1,5 million cells/ml for one week. In both cases filtered (0.2 μm), degassed, supernatant was adjusted to pH=7 and loaded onto a High-Trap Prot G column (GE Healthcare, Chicago, Illinois, USA) using an Äkta system (GE Healthcare, Chicago, Illinois, USA). Antibodies were eluted with a pH gradient ranging from pH 5.0-2.7 using 0.1 M trisodium citrate or 0.1 M glycine-HCL. 1000μl fractions were collected in tubes containing 175 μl Tris-HCl, pH=9 and dialyzed in dialysis cassettes from Thermo Fisher (Slide A_Lyzer Dialysis Cassette Cat. Nr. 87730) against PBS. Protein purity was assessed by SDS PAGE followed by Coomassie staining and Western blot analyses as described previously [57]. Protein concentration was determined by Bradford assay (Pierce, Rockford, USA).

### Virus propagation

The SARS-CoV-2 strain MUC-IMB-1 (GISAID EPI ISL 406862 Germany/BavPat1/2020) was isolated from an early outbreak cluster in Bavaria [39, 40] and was propagated by infection of Vero-E6 cells (DSMZ, Braunschweig, Germany) in DMEM (Gibco, ThermoFisher Scientific, Waltham, USA) supplemented with 10% heat-inactivated fetal calf serum (FCS, Capricorn Scientific GmbH, Ebsdorfergrund, Germany), 1% penicillin/streptomycin (Gibco, ThermoFisher Scientific, Waltham, USA) and 2 mM L-glutamine (Gibco, ThermoFisher Scientific, Waltham, USA). A second virus isolate (hCoV-19/Germany/ER1/2020; CoV-ER1) was obtained from a COVID-19 patient in Erlangen, passaged twice on Vero-E6 cells in OptiPRO™ (ThermoFisher Scientific, Waltham, USA) and sequenced (GISAID: EPI_ISL_610249). Supernatants of passaged viruses were harvested and filtered through a 0.45 μm cellulose acetate membrane filter. Aliquots of the supernatant were stored at −80°C until further use. The SARS-CoV-2 B.1.1.7 (GISAID EPI ISL 755639) and B.1.351 [62] variants of concern were obtained from a patient in München or Frankfurt, respectively. Following they were propagated as described previously for the CoV-ER1 variant.

The infectious titers were determined by limiting dilution in 96-well plates. Three days after infection, culture supernatants were removed from the wells and cells were washed with PBS and fixed with 4% paraformaldehyde in PBS for 20 min. Afterward, they were permeabilized for 15 min with 0.5% TritonX in PBS and blocked with 5% skimmed milk diluted in PBS for 1 hour. Subsequently, they were stained with protein G purified sera from a convalescent patient diluted 1:100 in PBS containing 2% skimmed milk. After 1 hour the cells were washed, and a goat anti-human IgG FITC (Jackson ImmunoResearch, West Grove, USA #109-096-088) antibody was applied. After 1 hour and a washing step, positive wells were identified by a CTL-ELISPOT reader (Immunospot; CTL Europe GmbH, Bonn, Germany). The signal was analyzed with the ImmunoSpot^®^ fluoro-X™ suite (Cellular Technology Limited, Cleveland, USA) software, and TCID50s were calculated as described previously [63].

### Virus neutralization assay

Neutralizing activities of hybridoma supernatants, antibodies and sera were assessed in a microneutralization assay based on detecting the number of virus-producing cells via immunofluorescence. For this, 1×10^4^ Vero-E6 cells were seeded per well of a 96-flat bottom plate one day before the infection. Purified antibodies, sera or hybridoma supernatants were first diluted and pre-incubated for one hour with 1,88×10E^4^ infectious units of SARS-CoV-2 stocks per well in a final volume of 100 μl. All dilutions of hybridoma supernatants or purified antibodies were prepared in cell OptiPRO™ culture medium. The dilution provided specifies the end-concentration of the antibodies after mixing with the diluted virus stock. After removing the cell culture medium, seeded Vero-E6 cells were incubated for one hour with 100 μl per well of the pre-incubated virus-antibody mix. Afterwards, the supernatant was discarded, the cells washed once with 100 μl PBS and 100 μl fresh cell culture medium added. The cells were incubated for 20-24 hours, washed with PBS, fixed with 4% paraformaldehyde, and subsequently stained as described above. Plates treated with murine antibodies were stained with protein G purified sera from a convalescent patient and plates treated with human antibodies with an antibody mix directed against SARS-CoV-2 at a concentration of 250ng/ml. IC50 values were calculated by plotting the virus activity in percent against the antibody concentrations and using the normalized response vs. inhibitor equation (variable slope) or inhibitor vs. response—Variable slope (four parameters) of GraphPad Prism 7.02.

### Quantitative antibody competition assay

For the assessment of the quantitative antibody competition SARS-CoV-2-S DNA transfected HEK-293T cells were incubated with 100 μl of a humanized protein G purified TRES antibody at a concentration of 250 ng/ml and serial dilutions of TRES antibodies with a murine Fc region at final concentrations ranging from 2.5 μg/ml-0.01 ng/ml. The cells were incubated for 30min on ice, washed, and bound antibodies were detected with a mouse IgG2a Alexa647-conjugated antibody directed against human IgG-Fc (BioLegend, San Diego, USA #409320). After washing the mean fluorescence intensities of transfected cells were determined with an Attune Nxt (Thermo Fisher Scientific, Waltham, USA) and the Flow Logic software™ (IInivai Technologies, Mentone, Victoria, Australia).

### Generation of escape mutants

5×10^6^ Vero-E6 cells were seeded on the day before infection in T175 flasks. TRES6 and TRES328 antibodies were incubated for one hour at a concentration of 200 ng/ml with 2×10^6^ TCID50 of the CoV2-ER1 virus. Subsequently, the medium in the cell culture flasks was exchanged for 20 ml of fresh medium and the virus/antibody mix was added to the cell cultures. Control cultures were inoculated with 2×10^6^ TCID50 of the CoV2-ER1 virus in the absence of any antibody. The cells were checked daily for infection. If infection occurred the C_T_ value was assessed by real time PCR. In case of a break through infection 100 μl of the break through viral supernatant were incubated with the antibody from the previous round at twice its concentration and added onto fresh Vero-E6 cells. This procedure was repeated five times. The TCID50 of the passaged virus stocks were measured and the neutralization against both TRES antibodies was assessed. Additionally the viral RNA was isolated from cell culture supernatant with a PureLink RNA Mini Kit (Thermo Fisher Scientific, Waltham, USA). The viral RNA was then sequenced for the identification of the escape mutations. Briefly, commercial reagents were used for unique dual indexed amplicon library generation according to the Artic protocol (NEBNext^®^ ARTIC SARS-CoV-2 Library Prep Kit (Illumina^®^)), which were sequenced with MiSeq^®^ Reagent Kit v2 (500 Cycles) on the Illumina MiSeq instrument and data analyzed with CLC Genomic Workbench 21 (Qiagen, Aarhus, Denmark). Median coverage obained for SARS-CoV-2^P5^ 1914, for SARS-CoV-2^TRES328^ 1326 and for SARS-CoV-2^TRES6^ 1436.

### Small animal challenge experiments

Animal experiments were performed following the EU Directive 2010/63/EU for animal experiments and were approved by local authorities after review by an ethical commission (TVV 21/20). Thirty-six female K18-hACE2 mice (Jackson Laboratory, Bar Harbor, USA) were infected intranasally under isoflurane anesthesia with 300 FFU of SARS-CoV-2 strain MUC-IMB-1 p.1 in a total volume of 50 μl. 24 hours after virus inoculation, twelve mice per group were injected intravenously either with 5.25 mg/kg body weight of TRES6 or TRES328 antibody or an isotype control antibody (TRES480) in a total volume of 100 μl. The animals were scored daily and the survival and disease incidence were measured over a maximal 10-day period. One cohort (n=6) was euthanized at day 4, while the second cohort (n=6) was scored for humane endpoints and euthanized at day 10 post virus inoculation. After euthanasia, BALs (bronchoalveolar lavages) were collected, and lung, brain, liver and spleen were taken. The organs were homogenized in 2 ml PBS using gentleMACS M tubes and gentleMACS Octo Dissociator (Miltenyi Biotec, Bergisch Gladbach, Germany). Afterward, the tubes were centrifuged at 2000 x g for 5 min at 4°C to discard cell debris. Viral RNA was isolated from 140 μl of homogenated supernatants or BAL using QIAamp Viral RNA Mini Kit (Qiagen, Aarhus, Denmark). The viral load in indicated organs and BAL was analyzed by RT-qPCR [64]. Reactions were performed using TaqMan^®^ Fast Virus 1-Step Master Mix (Thermo Fisher Scientific, Waltham, USA) and 5 μl of isolated RNA as a template. Synthetic SARS-CoV-2 RNA (Twist Bioscience, San Francisco, USA) was used as a quantitative standard to obtain viral copy numbers. The viral load reduction was calculated in comparison to the isotype control. The clinical scoring system included the following items: weight loss and body posture (0–20 points), general conditions including the appearance of fur and eye closure (0–20 points), reduced activity and general behavior changes (0–20 points), and limb paralysis (0–20 points). Mice were euthanized when reaching a cumulative score of 20.

For the detection of infectious virus in BAL Vero-E6 cells were seeded at 2×10^4^ cells/well in a 96-well plate in 200 μl of DMEM, 10 % FCS, 1× penicillin/streptomycin (Thermo Fisher Scientific, Waltham, USA) 24 h before infection with 20 - 100 μl of BAL diluted in DMEM with 1x penicillin/streptomycin for 3 hours. After replacing the supernatant with overlay medium (DMEM with 1 % methylcellulose, 2 % FCS and 1x penicillin/streptomycin), cells were incubated for 27 hours. SARS-CoV-2 infected cells were visualized using SARS-CoV-2 S protein specific immunochemistry staining with anti-SARS-CoV-2 spike glycoprotein S1 antibody (Abcam, Cambridge, Great Britain) as described previously [64]. Statistical evaluation of the data was performed by Kruskal-Wallis test (one-way ANOVA) and Dunn’s Pairwise Multiple Comparison Procedures as post hoc test.

## AKNOWLEDGEMENTS

We would like to thank Isabell Schulz and Doris Jungnickl for her excellent technical assistance. We kindly thank Jasmin Fertey and Rosina Ehmann for providing high titer virus stocks of SARS-CoV-2 and Alexandra Rockstroh for the optimized SARS-CoV-2 detection protocol. The expression plasmid for the spike ectodomain was kindly provided by Jason McLellan, Austin, USA, the TriGrid electrode array for DNA electroporation by Drew Hannaman, Ichor Medical Systems, Inc. and S2 and S1 protein by Thomas Schumacher, Virion/Serion GmbH, Würzburg. The ELISPOT Analyzer was obtained with financial support from Fondation Dormeur, Vaduz.

This work was supported by a grant (01KI2043) from the Bundesmininisterium für Bildung und Forschung (BMBF) and funds from the Bavarian State minstery for Science and the Arts to K.Ü., H-M J., and T.H. W. Further support was provided by B-FAST and COVIM, two BMBF-funded projects of the Netzwerk Universitätsmedizin (NaFoUniMedCovid19; FKZ: 01KX2021) and a DFG-funded research training group (RTG 2504).

The authors declare that they complied with all relevant ethical regulations. All relevant data supporting the findings of this study are available within the paper and its supplementary information files.

