## Supplement for "A pair of non-competing neutralizing human monoclonal antibodies protecting from disease in a SARS-CoV-2 infection model"

**Supplemental Material and Methods, Peter et al.,**

**Cell-cell fusion assay.** A cell-cell fusion assay was performed with HEK-293T cells transiently transfected with the SARS-CoV-2-S DNA and Vero-E6 cells constitutively expressing ACE2. For this $10^7$ HEK-293T cells were transiently transfected with SARS-CoV-2-S DNA and a blue fluorescent protein (BFP) expression plasmid by standard PEI transfection. The following day $10^7$ Vero E6 cells were stained with CellTraceTM CFSE (Thermo Fisher Scientific, Waltham, USA) according to the manufacturer recommendations, and plated into a flat bottom 96 well-plate. 48h post transfection the transfected HEK-293T cells were detached by washing and incubated with the diluted antibodies prior to addition to the CFSE labelled Vero-E6 cells at a ratio of 1:1. The cells were co-incubated for 45min and then trypsinized, washed with PBS and fixed with 2% PFA for 20 min at RT. The cells were analyzed on an FACS Attune Nxt and the percentage of CFSE and BFP double positive cells was determined using FlowJoTM (Treestar, Ashland, USA) software. The percent reduction of the percentage of double positive cells by the TRES antibodies was calculated by dividing the percentage of double positive cells for each antibody dilution by the percentage of double positive cells in the absence of the antibody. The IC50 was calculated by application of a 4-parameter curve fit using GraphPad Prism 7.02.

**Detection of TRES and TREShu antibody binding against Spike variants**

HEK-293T cells were transfected with plasmids encoding HA-tagged S proteins of the D614G variant of the B.1 strain, the B.1.1.7 variant or the B.1.351 variant. 48 hours after transfection the cells were incubated for 30 min with of TREShu antibodies at a concentration of 1000 ng/ml in FACS buffer and after washing, stained with a secondary antibody directed against human Fc (Biolegend, San Diego, USA, #409320). Following the cells were washed and fixed for 20 min with 2% PFA in PBS and then permeabilized with saponin (0.5% in FACS Buffer) for 10 min. Thereafter the C-terminal HA tag was detected with an anti-HA FITC labelled antibody (Sigma Aldrich, Taufkirchen, Germany, #7411) diluted in FACS buffer with 0,5% saponin. After Flow Cytometry the binding indices were determined: Binding index = (% TREShu positive cells * MFI of TREShu positive cells)/(% HA positive cells * MFI of HA positive cells).

**Supplemental Figures, Peter et al.,**

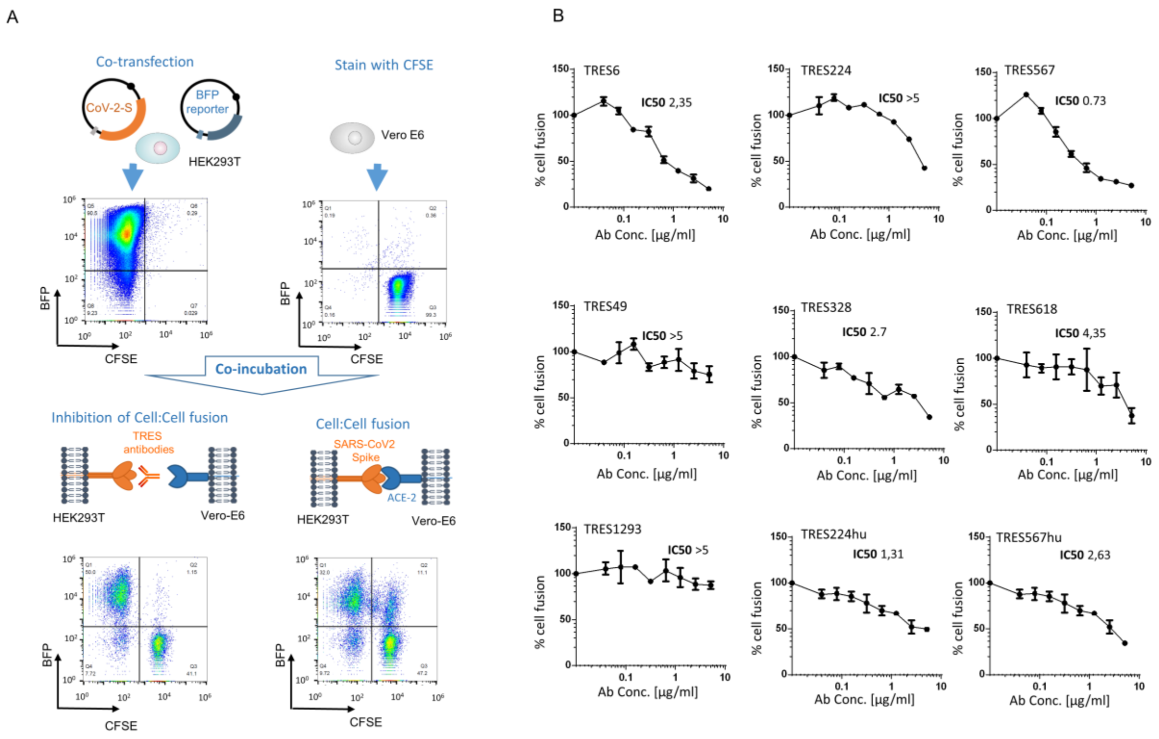


**Figure S1: Cell-cell fusion inhibition by addition of spike binding antibodies (A)** HEK-293T cells were transiently co-transfected with a BFP reporter plasmid and an expression vector encoding the complete spike protein. One day later Vero-E6 cells, constitutively expressing ACE2 were stained with CFSE. 48 h post transfection and 24h post labeling Vero E6 and HEK-293T cells were mixed and incubated 1:1 for 45 min in presence of different dilutions of spike binding antibodies. Double positive cells were identified by gating on Vero-E6 cells and the BFP+ CFSE+ cells. **(B)** The percentage of CFSE BFP double positive cells was acquired and the fusion inhibition determined by dividing the percentage of double positive cells for each antibody dilution with the percentage of double positive cells non-incubated with antibodies. The IC50 was calculated in µg/ml by application of a 4-parameter curve fit using GraphPad Prism 7.02. One representative experiment, out of two performed in duplicates is shown.

### TRES Cluster 1 sequences

```
                                   CDR1                       CDR2                                              CDR3
VH-GL  QVQLVESGGGVVQPGRSLRLSCAASGFTFSSYGMHWVRQAPGKGLEWVAVIWYDGSNKYYADSVKGRFTISRDNSKNTLYLQMNSLRAEDTAVYYCARETVDGMDVWGKGTTVTVSS
VH567  ...V...................................................................................V........V...........Q........
VH6    ...V...................................................................................V........V...........Q........
VH224  ...V..........................G..........................Q.............................V......H.V...........Q........
```

```
                                 CDR1                      CDR2                                         CDR3
Vκ-GL  NIQMTQSPSAMSASVGDRVTITCRARQGISNYLAWFQQKPGKVPKHLIYAASSLQSGVPSRFSGSGSGTEFTLTISSLQPEDFATYYCLQHNSYPCSFGQGTKLEIK
Vκ567  .................S.........D.N........................L........................................YT..........
Vκ6    .................S.........D.N........................L........................................YT..........
Vκ224  .................S.........D.N........................L........................................YT..........
```

### TRES Cluster 2 sequences

```
                                    CDR1                       CDR2                                                        CDR3
VH-GL   QVQLVQSGAEVKKPGASVKVSCKVSGYTLTELSMHWVRQAPGKGLEWMGGFDPEDGETIYAQKFQGRVTMTEDTSTDTAYMELSSLRSEDTAVYYCATAPAVAGPFYYYYYGMDVWGQGTTVTVSS
VH618   ...............................V.V.....................A......................................F............NF...I.............
VH1209  ...............................V.V.......E.............A......................................F...............................
VH1293  ...............................V.V.....................AK.............................S.......F...............................
VH49    ...............................V.V.....................A...............................Y......F...............................
VH328   ..H..........................V.V.V..........F.........NAA..............................Y......F..........L....................
VH219   ...........................S.I.V.V....................NA.....RG.R.....................KY......F...............................
```

```
                                      CDR1                        CDR2                                          CDR3
Vκ-GL    DIVMTQTPLSSPVTLGQPASISFRSSQSLVHSDGNTYLSWLQQRPGQPPRLLIYKVSNRFSGVPDRFSGSGAGTDFTLKISRVEAEDVGVYYCTQATQFPHSFGQGTKLEIK
Vκ618    ......................C..............................................................................T..........
Vκ1293   ......................C..............................................................................T..........
Vκ1209   ......................C............................I........C........................................T..........
Vκ49     ......................C...................................................E..........................T..........
Vκ328    ......................C...................................................E..........................T..........
Vκ219    ......................C...................................................E..........................T..........
```

**Figure S2: Alignments of the mature V regions of neutralizing TRES cluster 1 and cluster 2 antibodies.** The virtual germline sequence is shown above the TRES sequences. Sequence annotations and positions of CDR regions were determined with the abYsis software program. Amino acids are shown in the one-letter code. The positions of CDRs using the Kabat (blue underlined) and IMTG (blue bold letters) algorithms are indicated. Identical Vk sequences in cluster 2 antibodies are presented in the same color. Periods, identical amino acids; CDR, complementarity determining region.

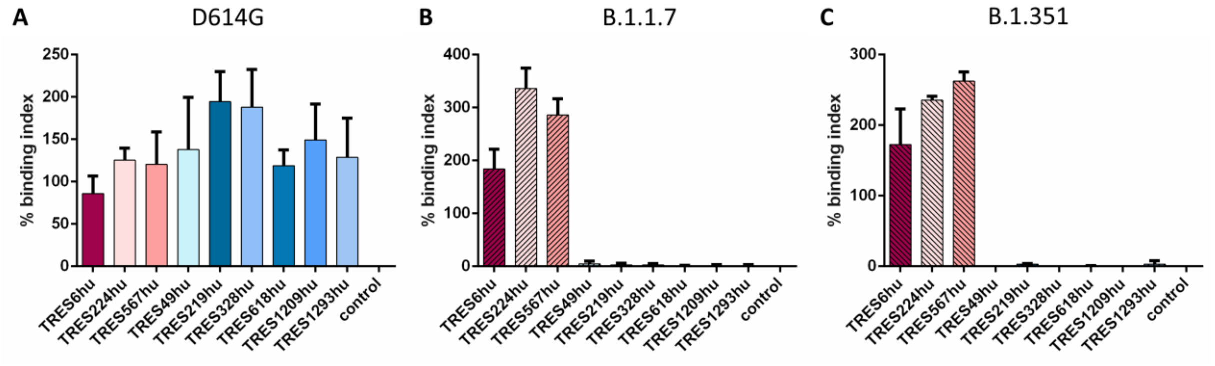


**Figure S3: Determination of binding indices against spike variants.** HEK-293T cells were transfected with plasmids encoding HA-tagged S proteins of the D614G mutant of the B.1 variant **(A)**, the B.1.1.7 variant **(B)** or the B.1.351 variant **(C)**. Cells were subsequently incubated with 1000 ng/ml of TRESHu antibodies. TRESHu antibodies bound to the spike protein were detected with an anti-IgGFc antibody. Additionally, the cells were stained for intracellular HA expression. The binding indices were calculated as described in Supplemental Material and Methods. One experiment with standard deviations performed in triplicates is shown.

**Supplemental Tables, Peter et al.,**

**Table S1: List of sequences of oligonucleotides**

| Name | Sequence | Description |
|---|---|---|
| Template Switch Oligo | GCTAATCATTGCAAGCAGTGGTATC AACGCAGAGTACATrGrGrG | TSO for 5'RACE RT |
| TSO Primer | CATTGCAAGCAGTGGTATCAAC | PCR Forward Primer |
| p350_mIgG1 | ATGGAGTTAGTTTGGGCAGCAGAT | Gene Specific PCR Reverse Primers |
| p354_mIgG2b | AGGAACCAGTTGTATCTCCACACC | |
| p616_mIgG2c | GAGCCAGTTGTACCTCCACACAC | |
| p355_mKappa | CTCCAGATGTTAACTGCTCACTGG | |
| p357_mLambda1 | ATCTACCTTCCAGTCCACTGTCAC | |
| p358_mLambda2/3 | ATTTGCCTTCCAGGCCACTGTCAC | |
| mCgamma1/2b/2c _Seq | GGCCAGTGGATAGACHGATG | mIgG PCR Sequencing Primer [1] |
| mKappa_Seq | CACTGGATGGTGGGAAGATGGATA | mKappa PCR Sequencing Primer [2] |


1. Bürckert, J.-P., et al., *High-throughput sequencing of murine immunoglobulin heavy chain repertoires using single side unique molecular identifiers on an Ion Torrent PGM.* Oncotarget, 2018. **9**(54): p. 30225-30239.

2. Chen, Y., et al., *Barcoded sequencing workflow for high throughput digitization of hybridoma antibody variable domain sequences.* Journal of Immunological Methods, 2018. **455**: p. 88-94.
